# Beta bursts in the parkinsonian cortico-basal ganglia network form spatially discrete ensembles

**DOI:** 10.1101/2024.03.05.583301

**Authors:** Isaac Grennan, Nicolas Mallet, Peter J. Magill, Hayriye Cagnan, Andrew Sharott

## Abstract

Defining spatial synchronization of pathological beta oscillations is important, given that many theories linking them to parkinsonian symptoms propose a reduction in the dimensionality of the coding space within and/or across cortico-basal ganglia structures. Such spatial synchronization could arise from a single process, with widespread entrainment of neurons to the same oscillation. Alternatively, the partially segregated structure of cortico-basal ganglia loops could provide a substrate for multiple ensembles that are independently synchronized at beta frequencies. Addressing this question requires an analytical approach that identifies *groups* of signals with a statistical tendency for beta synchronisation, which is unachievable using standard pairwise measures. Here, we utilized such an approach on multichannel recordings of background unit activity (BUA) in the external globus pallidus (GP) and subthalamic nucleus (STN) in parkinsonian rats. We employed an adapted version of a principle and independent component analysis-based method commonly used to define assemblies of single neurons (i.e., neurons that are synchronized over short timescales). This analysis enabled us to define whether changes in the power of beta oscillations in local ensembles of neurons (i.e., the BUA recorded from single contacts) consistently covaried over time, forming a “beta ensemble”. Multiple beta ensembles were often present in single recordings and could span brain structures. Membership of a beta ensemble predicted significantly higher levels of short latency (<5ms) synchrony in the raw BUA signal and phase synchronization with cortical beta oscillations, suggesting that they comprised clusters of neurons that are functionally connected at multiple levels, despite sometimes being non-contiguous in space. Overall, these findings suggest that beta oscillations do not comprise a single synchronization process, but rather multiple independent activities that can bind both spatially contiguous and non-contiguous pools of neurons within and across structures. As previously proposed, such ensembles provide a substrate for beta oscillations to constrain the coding space of cortico-basal ganglia circuits.

## Introduction

Beta oscillations (15 to 35 Hz) across the primary motor cortex and basal ganglia are enhanced during Parkinson’s disease (PD) (Brown, 2007; Hammond et al., 2007). Effective therapies for PD, such as deep brain stimulation (DBS) and levodopa administration, reduce beta power and the degree of suppression correlates with improvement in bradykinetic symptoms (Brown et al., 2001; Kuhn et al., 2008). Beta oscillations are not stationary, but rather occur in “bursts” that consist of transient increases in the instantaneous power of basal ganglia local field potentials (LFPs) or the electrocorticogram (ECoG) (Feingold et al., 2015; Cagnan et al., 2019). OFF levodopa, beta bursts increase in duration and amplitude (Tinkhauser et al., 2017a). The reduction in the number of beta bursts and suppression of long-duration bursts by treatment (e.g. levodopa or DBS) are correlated with the degree of improvement in motor performance (Little et al., 2013; Tinkhauser et al., 2017a).

A key question in the field of PD pathophysiology is why enhanced beta oscillations lead to motor symptoms. Several authors have proposed that the level of beta synchronisation influences the dimensionality of the coding space in the cortico-basal ganglia circuits (Engel et al., 2001; Bar-Gad et al., 2003; Brittain et al., 2014). In this framework, individual neurons or small ensembles of neurons should have a relatively large degree of independence to function optimally. This is supported by the low-level of temporal correlation between basal ganglia neurons in awake animals (Raz et al., 2001). Short periods of enhanced synchronisation (<1s), however, could be utilised to hold a population of neurons in a given state (Engel et al., 2001; Brittain et al., 2014). In the healthy motor system, transient periods of synchronisation could support muscle synergies to hold part of the limb still (Bracklein et al., 2022). However, if these periods of synchronisation become abnormally sustained, they could impair dynamic processing and execution of the associated behaviour.

Beta synchronisation can occur in both temporal and spatial dimensions. Detailed examination of beta bursts has demonstrated that beta oscillations in the parkinsonian brain are pathologically extended in time (Tinkhauser et al., 2017b; Tinkhauser et al., 2017a). The length of the cortical (ECoG) burst predicts duration of cortico-basal phase-locking at the level of single– and multi-units, suggesting that the elongation of beta bursts in LFP is representative of oscillation at the level of neuronal output (Cagnan et al., 2019). Given that field signals (EEG/ECoG/LFP) are the result of temporally synchronised synaptic potentials across neurons (Buzsaki et al., 2012), sustained bursts also indicate enhanced spatial synchronisation at a local level around the electrode. However, any spatial synchronisation between different field potentials recorded within or across structures, indicating coupling over larger spatial scales, will be a composite of volume conduction, the reference configuration and physiological coupling. LFP/ECoG-based analyses will thus provide only superficial insights into spatial synchronisation at the level of neuronal populations.

With respect to unit activity, the beta coherence between the spiking of pairs of units in the rodent GP decreases with distance between the recorded pair (Mallet et al., 2008b). However, neurons in the cortex, basal ganglia and thalamus are clustered into hundreds of semi-independent pathways that may not conform to simple gradients in one or more anatomical plane (Hunnicutt et al., 2016; Foster et al., 2021). Neurons in a given brain structure could divide bimodally into populations that always do or do not engage with the same unified process. Alternatively, there could be independent assemblies of neurons that are synchronised at beta frequencies at different times. To distinguish between these scenarios requires analyses of the spatiotemporal dependence of the synchronisation of neuronal output using a method that does not rely on signals being spatially contiguous and that can detect different ensembles of neurons that are independent of one another.

Here we employed an analytical framework using principal component and independent component analyses, commonly used to define groups of neurons with a tendency to co-fire over short-timescales (i.e. neural ‘ensembles’). We applied this approach to multichannel background unit activity (BUA) recorded in 6-OHDA-lesioned rats in the external globus pallidus (GP) and subthalamic nucleus (STN) to define features of the spatiotemporal synchronisation of pathological beta oscillations. The results show that multiple, independent ensembles of neurons are coordinated by the temporal dynamics of ongoing beta oscillations, and that neurons comprising these ‘beta ensembles’ can be distributed across brain structures.

## Methods

### Parkinsonian rats

This study uses data from Mallet et al, 2008. Extracellular recordings were made using multichannel silicon probes in the GP and STN of urethane-anaesthetised Sprague-Dawley rats (n=15) rendered parkinsonian by 6-OHDA-lesioning. 6-OHDA lesioning was carried out under anaesthesia by injecting 1 microL of 6-OHDA 1,2-1,4 mm lateral and 4.1 mm posterior to Bregma and 7.9mm ventral to the dura. Animals that recovered successfully and were deemed to have been effectively lesioned were used for electrophysiological recordings. Lesions were classified as effective if 15 days after lesioning on administration of apomorphine animals performed >= 90 contraversive rotations in 20 minutes. Recordings were made on a pair of probes, each with 16 electrodes arranged in a single vertical plane separated by 100 microns. In some recordings, one probe was in the GP and one in the STN (n = 7/45 recordings) whereas, in others, probes made up of two shanks separated by 500 microns were targeted to the GP (n = 38/45 recordings). The number of electrodes within targeted structures varied between recordings. Channels outside of the STN and GP were excluded from the analysis. Frontal ECoG recordings, ipsilateral to the lesion, were made simultaneously with those in the basal ganglia using a 1mm-diameter screw above the frontal motor cortex referenced to a screw overlying the ipsilateral cerebellar hemisphere. Signals were recorded using a Power 1401 amplifier and Spike2 (Cambridge Electronic Design Limited) at a sample rate of 16 kHz. Following recordings animals were euthanised and fixed by transcranial perfusion.

### Signal processing of background unit activity (BUA) and beta envelope

Background unit activity (BUA) was derived from raw probe recordings in line with several previous studies (Moran and Bar-Gad, 2010; Sharott et al., 2017; Cagnan et al., 2019; Nakamura et al., 2021). First, we high-pass filtered at 300Hz using a 3^rd^ order Butterworth filter. Large spikes that could dominate the signal were identified by setting a threshold of mean ± 4 SD of the recording, and then a 4 ms segment around each instance crossing this threshold was removed and replaced with a randomly selected epoch which did not contain any spiking activity (Cagnan et al., 2019). Data was rectified and then low-pass filtered at 300 Hz using a third order Butterworth filter. BUA was down sampled to 1000Hz for subsequent analysis. BUA was then filtered around ± 5 Hz of the frequency of maximum coherence with the frontal ipsilateral ECoG in the beta frequency range (15-35Hz), using a 2^nd^ order Butterworth filter. The instantaneous envelope of beta oscillations was computed from the magnitude of the Hilbert transform of the beta filtered BUA. The change in the envelope of beta oscillations over 50ms was calculated for each envelope-timeseries. The first and last 200ms of data were removed to avoid any edge effects. On average, 2153.9 ± 894 data points (differenced envelope of beta filtered BUA) were computed per recording over the 45 recordings. All filtering was conducted using zero phase lag filters.

### Beta ensemble identification

Beta ensembles were identified using a two-step statistical process, which allowed us to identify groups of channels with correlated changes in beta envelope (Lopes-dos-Santos et al., 2011; Lopes-dos-Santos et al., 2013; van de Ven et al., 2016).

#### Principle component analysis

A matrix *Z* was constructed where the z-scored change in the envelope of beta filtered BUA over 50ms for each channel made up a column. The matrix *Z* is, therefore, a *T* × *N*, where *T* is the number of 50ms intervals that the change in envelope of the beta filtered BUA is calculated over and *N* is the number of channels.

Principal component analysis (PCA) was then applied to the matrix *Z*. To determine the number of significant patterns embedded in the data, each column of matrix *Z* had a random number of consecutive elements shifted from its end to its start, giving rise to the matrix the *Z_shifts_*. The data for each channel in the rows of *Z_shifts_*, therefore, are no longer from the same point in time and any co-ordination detected arises due to chance. PCA was then performed on *Z_shifts_* and the resulting eigenvalues were stored. This process was repeated 1000 times to produce a distribution of eigenvalues under null conditions. A significance threshold was set using a Bonferroni correction for the number of principle components tested. Given that the number of principal components is equal to the number of channels included in analysis, 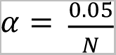. If an eigenvalue from PCA on *Z* was greater than the value of the distribution of eigenvalues produced from the shifted data at the 100(1-α)*th* percentile, the corresponding principal component was considered significant. Significant principal components were stored a *N* × *S* matrix, *P_sign_*, *S* is the number of significant principal components.

#### Independent component analysis

To avoid constraining assembly patterns to be orthogonal and to allow the detection of higher-order correlations, independent component analysis (ICA) is used (Lopes-dos-Santos et al., 2011; van de Ven et al., 2016). Data is first projected onto the subspace spanned by the significant principal components, followed by ICA to prevent the detection of spurious patterns. The unmixing matrix was expressed in the original basis. This results in a *N* × *S* matrix, where N is the number of channels and S is the number of significant principal components. The columns of this matrix are the weight vectors of the assembly pattern. The sign and size of these weights are arbitrary, so each column of V was scaled to unit length, and the sign of each column of V was set such that its largest vector weight was positive. The background unit recorded by a channel was said to be a member of an ensemble if its absolute weight in the assembly pattern was greater than the 80^th^ percentile of absolute weights.

### Eigenvalues and frequency

The relationship between the frequency band of BUA and the strength of the correlations identified by PCA were assessed by filtering BUA at ± 5Hz using a second order Butterworth filter around a centre frequency varying from 8Hz to 100Hz in 1Hz increments. The change in envelope was then computed and PCA was applied. The duration over which the change in envelope was computed was the period of the centre frequency, so the number of samples PCA was conducted on depended on the centre frequency of the oscillation. However, the magnitude of eigenvalues that arise due to chance changes as a function of the number of samples included in PCA. To account for this, eigenvalues were normalised by subtracting and then dividing by the significance threshold for ensembles (see above) for that frequency band. The normalised eigenvalues therefore reflect proportionally how much greater an eigenvalue is then would be expected under null conditions.

The magnitude of eigenvalues and number of significant ensembles were compared in the beta (15-25Hz), low-gamma (35-60Hz) and high-gamma (60-90Hz) frequency bands (filtering was carried out using a 2^nd^ order Butterworth filter). Again, because the change in envelope was calculated over the period of gamma oscillations, which is shorter than that of beta oscillations, there were more data points for both the low and high gamma band. To avoid variability in the magnitude of eigenvalues due to chance between frequency bands, changes in the envelope of gamma band oscillations were randomly subsampled without replacement to match the same number of data points as were available for beta oscillations. As a result, the same number of data points were subjected to PCA for each frequency band, making the eigenvalues comparable across conditions.

### Assembly expression strength

The expression strength of the assembly pattern was then calculated for each point in time:

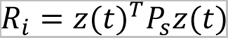

where *z*(*t*) is a column vector whose elements are the z-scored change in envelope of beta filtered BUA at time *t* for each of the channels and *P_s_* is the outer product of each column of *V* (s = 1, …, *S*):

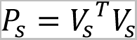

after the elements in the leading diagonal are set to 0:

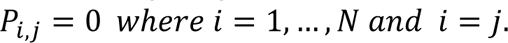

Setting the lead diagonal to 0 ensures that a change in the envelope in the beta filtered BUA in a single channel with a large contribution to the assembly pattern is not sufficient to bring about high ensemble expression-strength. A threshold of 5 was set to define an ensemble activation, as has previously been used for ensembles of single units (van de Ven et al., 2016, Trouche et al., 2016). These activations were used as a trigger point for averaging the envelope and phase synchrony of beta oscillations.

### Phase synchrony

The instantaneous phase of beta filtered BUA was derived from the Hilbert transform. Phase synchrony index (PSI) was then computed for overlapping 50 ms intervals for pairs of channels *a* and *b*:

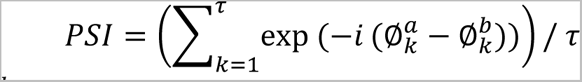

where 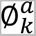 and 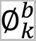 is the instantaneous phase of the beta filtered BUA of channel *a* and *b* respectively, and τ is the number of samples that make up the 50ms windows. While non-overlapping windows were used for PCA-ICA to avoid inflating the magnitude of eigenvalues, here an overlapping window allows for greater temporal resolution into the changes in phase synchrony.

### Correlations between signals

Correlations and cross-correlation between the change in beta envelope over 50ms bins in members and non-members were calculated across all permutations of pairs of channels and then averaged for each ensemble. Similarly, when calculating the correlation or cross-correlation at a given distance, all permutations of pairs of channels at that distance were computed and then averaged. Whether pairs were more positively correlated than expected due to chance (e.g., Fig 1F) was assessed by circularly shuffling the change in the envelope of the beta filtered BUA for each pair of channels 1000 times. If the true correlation exceeded the 95^th^ percentile for shuffled data a pair of channels was classified as significantly correlated.

**Figure 1.**
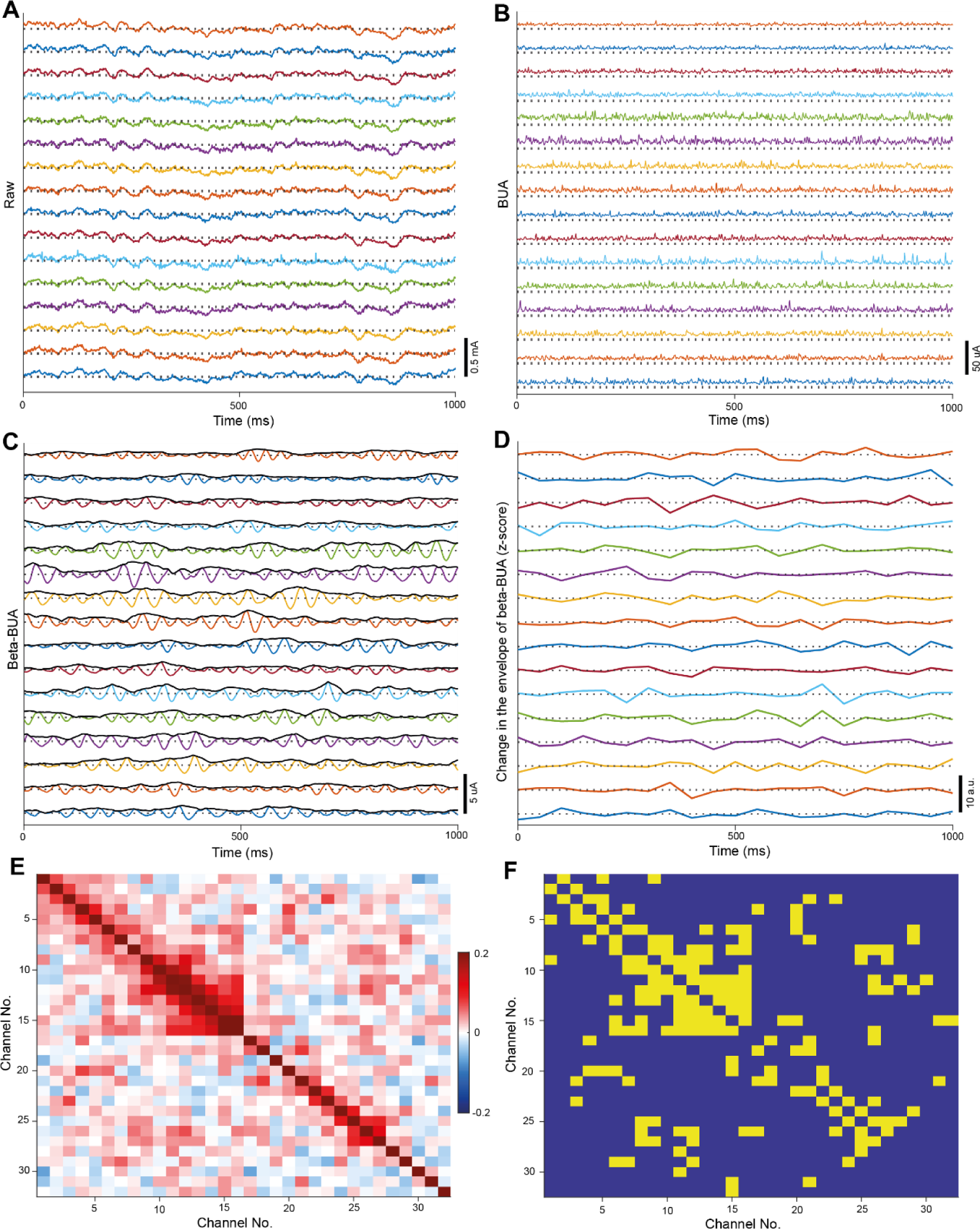
Analysing covariance in the change in envelope of beta-BUA within and across basal ganglia structures: Recordings were made with a pair of multi-electrode probes either both targeted to the GP or one targeted to the GP and the other to the STN. Either a single probe with two 16 channel shanks (targeted to the GP) or two probes each with 16 channels (targeted to the GP and STN) were used. Each shank in both configurations consisted of 16 electrodes arranged in a single vertical plane separated by 100 microns. Data is plotted here **A**: The wideband signal recorded from 16 channels on a single recording shank in the GP. The dotted line represents zero for each signal. **B**: The BUA from 16 channels on a single recording shank in GP. The dotted line represents zero for each signal. **C**: The beta filtered BUA (coloured) and the envelope of this signal (black) from 16 channels on a single recording shank in the GP. The dotted line represents zero for each signal. **D**: The z-scored change in envelope of the beta filtered BUA over 50ms from 16 channels on a single recording shank in the GP. The dotted line represents zero for each signal. **E**: An exemplary correlation matrix from a single recording session showing the correlations between the change in the envelope of beta filtered BUA over 50 ms windows for pairs of channels. The channels numbered 1-16 were from probe 1, whereas the channels numbered 17-32 were from probe 2. Channels are numbered by their position on the probe (e.g., channels 2 and 3 neighbour each other on probe 1). **F:** The significantly correlated pairs of channels (yellow) in the correlation matrix **E**. The main diagonal, which displays the correlation of a channel with itself, was displayed as non-significant in matrix **F** as significant correlations here are not meaningful.

### Analysis and plotting

Analysis and plotting were carried out using custom written code in MATLAB (Mathworks, Natick, MA, USA). Error bars represent standard error of the mean (SEM) unless otherwise stated.

## Results

Our overarching aim was to identify the spatial extent of basal ganglia unit activity that was coordinated by pathophysiological beta oscillations. Unlike LFPs, background unit activity (BUA), which reflects the synchronous spiking of the neural population proximal to the recording electrode, is a well spatially localised signal. Background unit activity (BUA) was computed from the GP and STN (**Fig 1A, B**) of anaesthetised, parkinsonian rats (Mallet et al., 2008a; Cagnan et al., 2014). A Hilbert transform was used to extract the envelope of the beta filtered BUA (**Fig 1C**), the change in the envelope over 50 ms windows was then computed (**Fig 1D**). This allowed us to detect changes in the amplitude of beta oscillations in the firing of neurons in the population of neurons proximal to the recording electrode. We focused on the correlations between the *change* in envelope across channels rather than the envelope itself, as it allowed us to identify groups of channels where the emergence and offset of beta oscillation occurred synchronously.

We examined the change in beta-envelope over 50ms as this is the approximate period of the beta oscillations observed here (centre frequency: 20.9Hz ± 2.7Hz). Moderate pairwise correlations were observed between the change in the envelope of beta filtered BUA (in channels across the GP and STN (**Fig 1E**). This was particularly the case for channels in close proximity, as indicated by the high correlations observed around the main diagonal of the correlation matrix. There were, also, several significant correlations observed over greater distances (**Fig 1F)**, including between channels on different recording probes. BUA is a far better localised signal, as compared with monopolar LFP, (**Fig S1**) so these correlations are unlikely to result from volume conduction. This demonstrates beta oscillations emerge in a coordinated fashion across channels, including channels that are not immediately adjacent or on the same recording probe.

### Synchrony and beta ensembles

We hypothesized that unit activity across the basal ganglia would be coordinated by *multiple* beta ensembles. To test this, we sought to identify *groups* of BUA signals recorded from different probe contacts (beta ensembles) that had co-ordinated changes in their oscillation strength. This was achieved using a two-step statistical process (Fig 2), PCA followed by independent component analysis (ICA), whereby *groups* of channels with correlated changes in beta envelope (**Fig. 2**) could be identified. 85 such significant beta ensembles were identified over 45 recordings in 15 animals, each described by a weight vector containing the contribution of each channel (**Fig. 2F**) to the beta ensemble.

**Figure 2.**
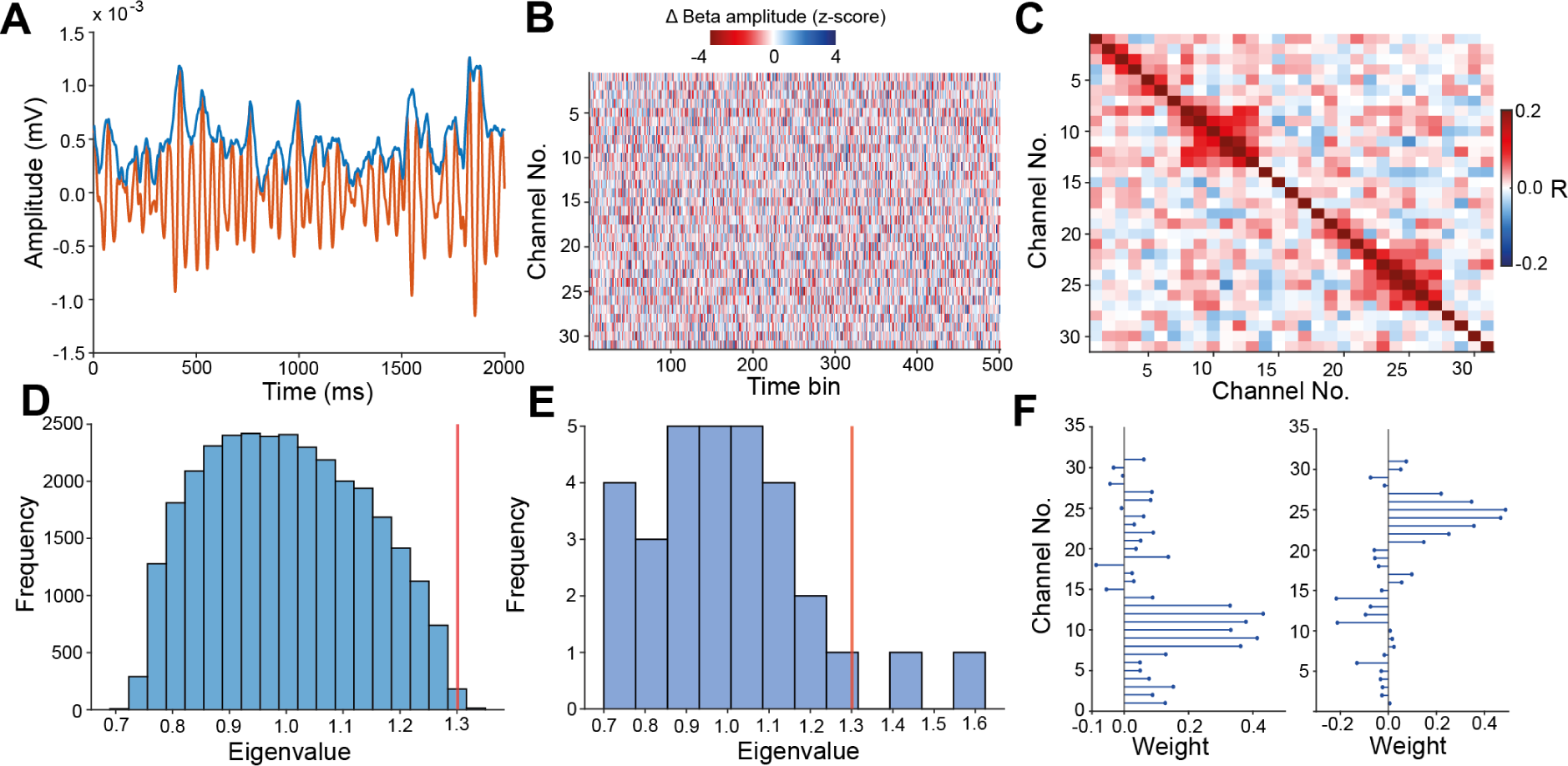
PCA-ICA was used to identify groups of channels with coordinated changes in the envelope of beta filtered BUA: All data presented here is from a single recording session in the GP. **A**: Exemplary beta-BUA (orange) and beta-envelope (blue) from a single probe channel in the GP. **B**: Exemplary z-scored change in beta-envelope over consecutive 50 ms windows for a single recording (only a section of the recording is shown here for display purposes). **C**: The correlation matrix of the z-scored change in beta envelope recorded over 31 channels in a single recording. All channels here were located in the GP. **D**: The distribution of eigenvalues from PCA applied to the matrix of z-scored changes in beta-envelope under null conditions (with randomly circularly shifted columns). The orange line shows the significance threshold determined by a percentile cut-off of the circularly shuffled data with Bonferroni correction. **E**: PCA was applied to the z-scored change in beta-envelope. Two principal components were classified as significant here on the basis that their eigenvalues were greater than the 95^th^ percentile (adjusted by the Bonferroni correction) of eigenvalues from the circularly shifted data (vertical red line). ICA was then applied to the z-scored change in the envelope of the beta filtered BUA projected onto the significant components to identify the assembly patterns of the 2 beta ensembles. **F**: The assembly patterns of 2 significant beta ensembles identified in this single recording session. Each recording channel has a weight in these assembly patterns which represents how large a contribution BUA recorded on that channel makes to the identified beta ensemble.

Channels with an absolute weight greater than the 80^th^ percentile of the weight vector were classified as members of the beta ensemble (**Fig 3**). In total, 85 beta ensembles were identified as significant on the basis of the magnitude of eigenvalues from PCA. In two recordings, the fastICA algorithm did not converge so these were excluded from analysis leaving 81 beta ensembles. Most recordings had probes only targeted to the GP (38/45) and as such 75 of the 81 ensembles were made up of channels in the GP alone. 6 of the 10 ensembles identified on the recording days with an STN recording probe contained at least 1 STN member, with 2 beta ensembles containing member channels both in the GP and STN. Member channels were generally clustered in space, although in some instances, member channels were separated by large distances (e.g., **Fig 3B, E and G**). The majority of ensembles (57/81) were made up of member channels that were spatially contiguous. However, some examples (13/81) of ensembles with non-spatially contiguous members within a single probe could be identified, as could examples of ensembles which spanned the two shanks of the recording probe (12/81).

**Figure 3:**
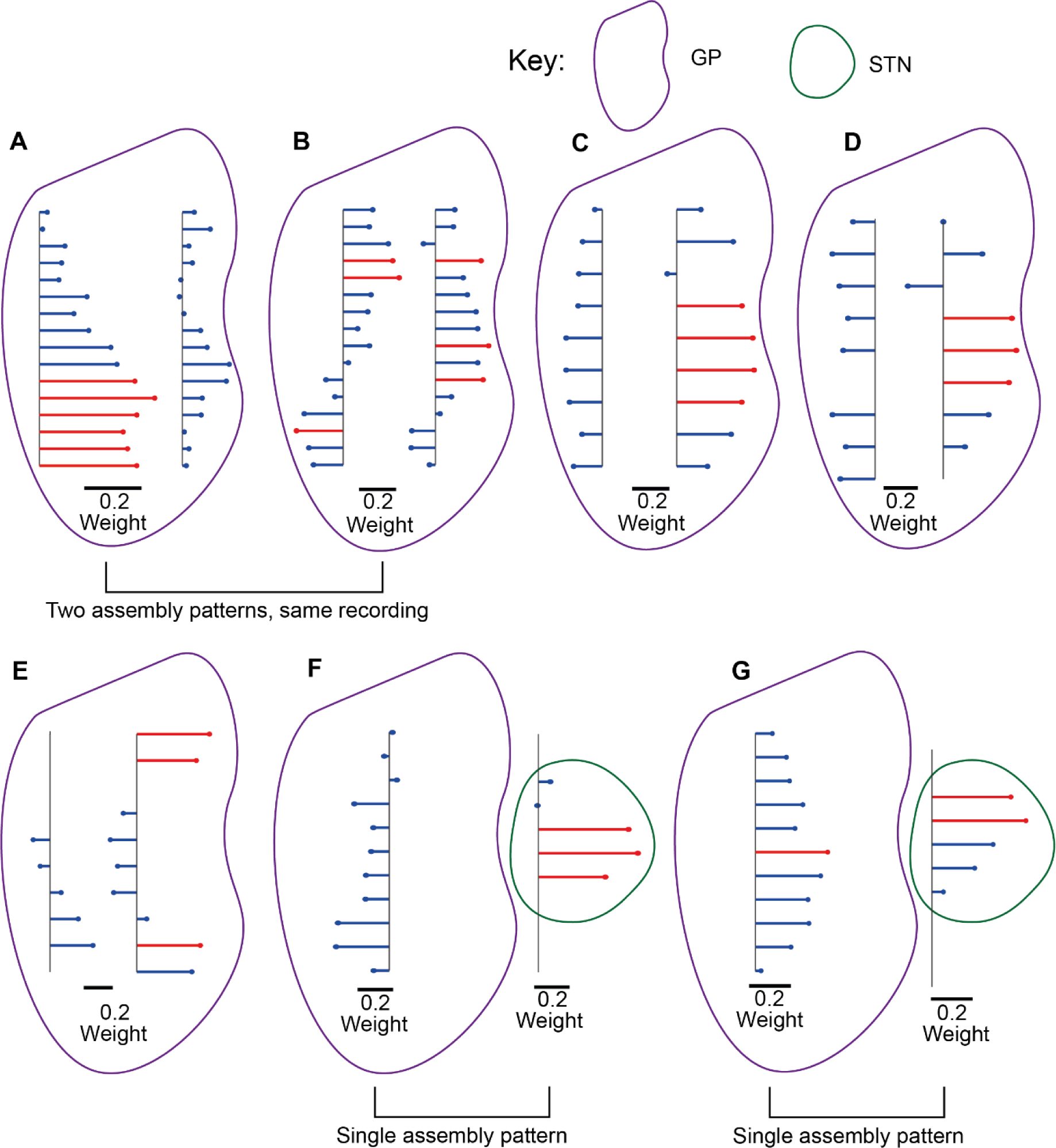
Beta ensembles were mostly spatially contiguous but could be non-contiguous, span shanks or even brain structures. **A-E**: Assembly patterns with a two-shank probe targeted to the GP. Member channels where coloured in red for display purposes. **F, G:** Assembly patterns with one probe targeted to the GP and another to the STN. Member channels where coloured in red for display purposes.

Strikingly, two beta ensembles were identified in most recordings (**Fig 4A**), confirming that beta oscillations are not a completely unified process, but occur in spatially segregated ensembles. The number of channels within the GP or STN varied considerably across recordings (**Fig 4B**), as did the number of member channels (**Fig 4C**). Ensembles most frequently had 2 or 3 member channels (**Fig 4C**) although some were made up of larger numbers (up to 6). There were some instances were only a single member channel could be identified for an ensemble. This was a result of a short falling of using a percentile-based cut-off to define assembly membership. In recordings with small numbers of channels (i.e., less than 10), an 80^th^ percentile cut-off did not consistently identify 2 or more sufficiently high weighted channels. None the less, these ensembles still represent a pattern of statistically significant coordination in the change in the envelope of beta filtered BUA. Member channels tended to be clustered spatially (**Fig 4D**), with most member channels being separated by 500 microns or less.

**Figure 4:**
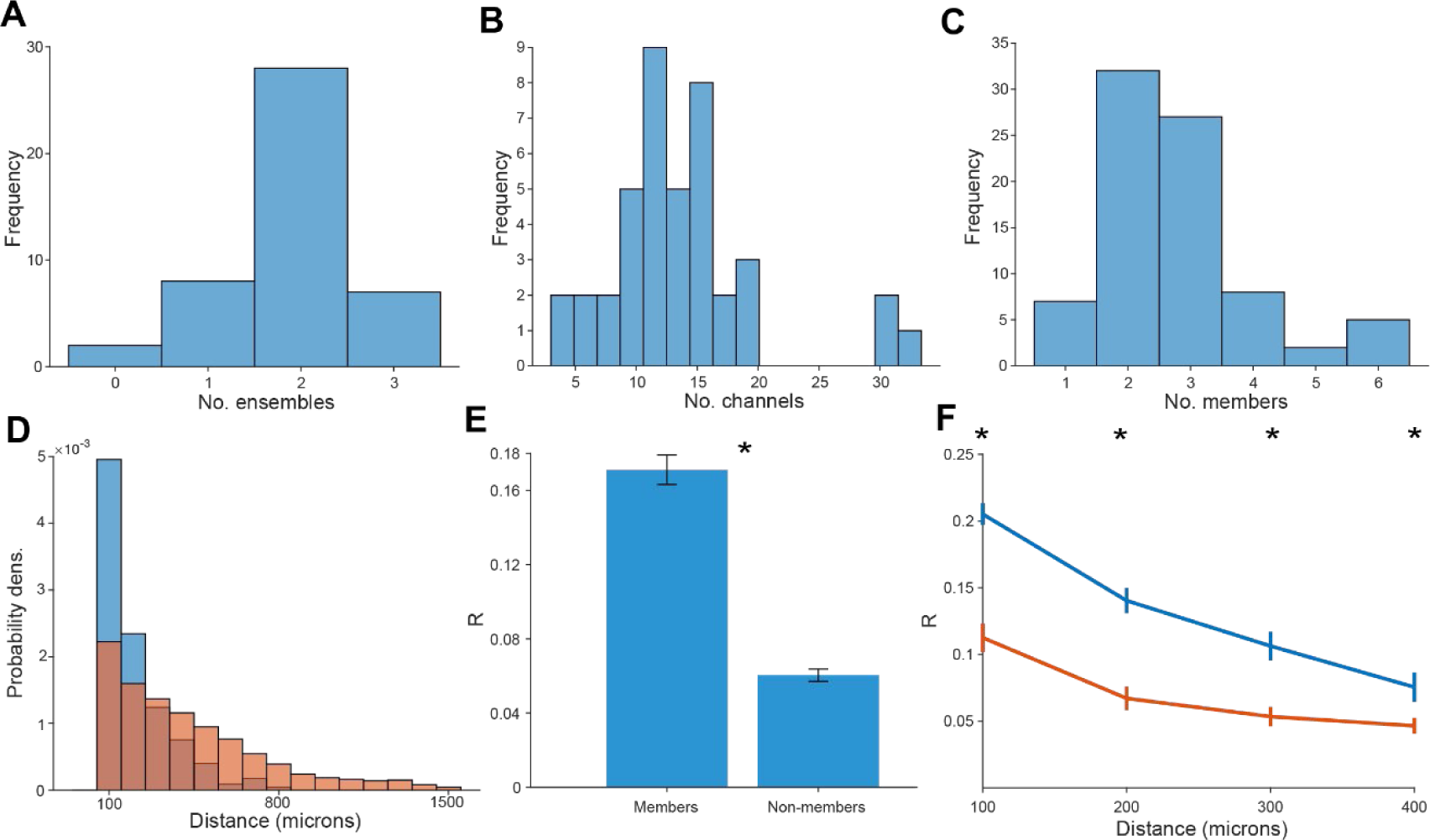
Beta ensembles were consistently identifiable across recordings and were made up of spatially clustered channels. **A**: A frequency histogram of the number of ensembles detected for each recording. **B**: A frequency histogram of the number of recording channels present in target structures (GP and STN) per recording. **C**: The frequency histogram of the number of member channels for each ensemble. **D**: The probability density histogram of the distance between pairs of member (blue) and non-member (red) channels on each recording probe individually. **E**: The average Pearson’s R for changes in the envelope of beta filtered BUA was significantly greater for pairs of member channels than for pairs of non-member channels (Wilcoxon signed-rank test, p=7.7 x 10^-14^). **F**: The average Pearson’s R was greater between pairs of member channels (blue) than pairs of non-member channels (orange), separated by 100, 200, 300 or 400 microns (Wilcoxon rank sum test, p<0.05 for 100, 200, 300 and 400 microns).

Post-hoc correlation of the change in envelope of beta filtered BUA was greater between pairs of member channels than pairs of non-member channels (**Fig 4E**), confirming that we had indeed identified channels with coordinated beta onset and offset. This was still the case when comparing correlations between channels separated the same distance (**Fig 4F**), so this effect is not solely driven by member channels being more proximal than non-members. Some highly weighted channels had a negative weighting in the assembly pattern, indicating that power changes in that channel were in the opposite direction to other channels. However, these instances were rare (9/224 channels that were assembly members) so were excluded from the analysis. Due to the relatively small number of STN recordings, we limited subsequent sessions where recordings were only made in the GP (71 beta ensembles) to simplify interpretation.

The member channels were identified in all analysis to this point were identified by changes in the *amplitude* of beta envelope. We hypothesised that *phase-synchrony* between members of beta ensembles would also be higher than non-members, which would indicate that they are also coordinated on the timescale of individual cycles. To address this, we explored cross-correlations between *unfiltered* BUAs calculated between ensemble members and non-members. If correlations in the beta power between channels predicted beta synchrony, beta fluctuations should emerge through summation, without the need to filter (see Cagnan et al, 2019). Cross-correlations between raw BUA showed a peak at 0ms lag and broader peaks at ±50ms (**Fig 5A**). The cross-correlation between members was greater than that between non-members at lags of 0ms and lags of multiples of 50ms corresponding to beta-frequency correlation (**Fig 5A)**, confirming that there is greater beta phase-synchronisation between member channels.

**Figure 5.**
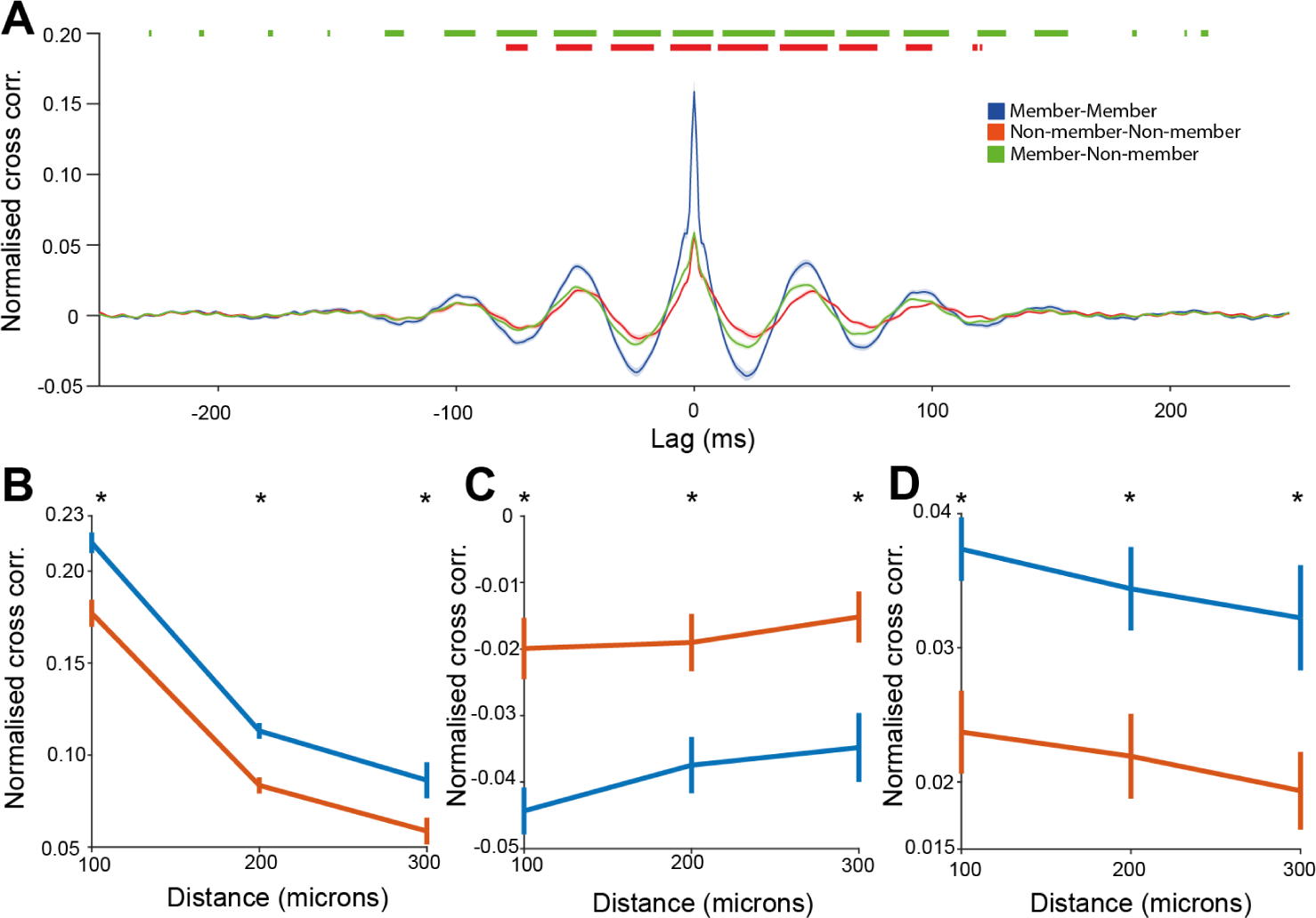
Members of beta ensembles showed correlated changes in the envelope of beta filtered BUA and increased beta band coherence. **A**: The normalized cross correlation between pairs of member channels (blue), pairs of non-member channels (orange) and pairs of member and non-member channels (green), at lags of –250ms to 250ms. Orange markers above the graph show significance between the cross correlation of pairs of members and pairs of non-member channels. Green markers above the graph show significance between the cross correlation of pairs of members and pairs of member and non-member channels. Significance was determined with the Wilcoxon rank sum test using false discovery rate statistics to control for the multiple time points compared. **B**: The normalized cross correlation at a 0ms lag between pairs of member channels was greater than that for pairs of non-member channels when separated by the distance between channels (Wilcoxon rank sum test, p<0.003 for 100, 200 and 300 microns). **C**: The normalized cross correlation at a 25ms lag between pairs of member channels was more negative than that of pairs of non-member channels when separated by the distance between channels (Wilcoxon rank sum test, p<0.004 for 100, 200 and 300 microns). **D**: The normalized cross correlation at a 50ms lag between pairs of member channels was greater than that for pairs of non-member channels when separated by the distance between channels (Wilcoxon rank sum test, p<0.006 for 100, 200 and 300 microns).

The cross correlation at 0ms in raw BUA between pairs of member and pairs of non-member channels fell with increasing distance **(Fig 5B)**, but was greater between member channels than between non-members even after taking distance into account **(Fig 5B)**. Equally, the trough at 25ms was lower **(Fig 5C)** and the peak at 50ms was higher **(Fig 5D)** in member channels as compared to non-members after controlling for the distance between channels. This indicates that differences in phase-synchrony between members and non-members is not simply a function of the average distance between channels.

### Beta ensembles were preferentially observed in the beta-band

Given that filtering is a key part of the signal processing pipeline, it is possible that the results described thus far was due to covariance in wideband, rather than beta band amplitude. To address, this we compared the core analytical variables to those computed on other frequency bands. The normalised variance explained by the first principal component was greater in the beta frequency range than in lower or higher frequency ranges (**Fig 6A**), with a peak at ∼20Hz. This was also true of the mean of the normalised variance explained by the first 3 principal components (**Fig 6B)**. Consistent with this finding, the eigenvalue of the first principle component (**Fig 6C)** and mean of the eigenvalues for the first three principal components (**Fig 6D**) was significantly greater for beta frequency than low or high gamma filtered BUA. In addition, significantly more beta ensembles could be identified than for either the low or high gamma band (**Fig 6E**). Therefore, oscillations at beta frequency led to more spatially coordinated neuronal activity than oscillations at higher frequency bands in parkinsonian animals.

**Figure 6:**
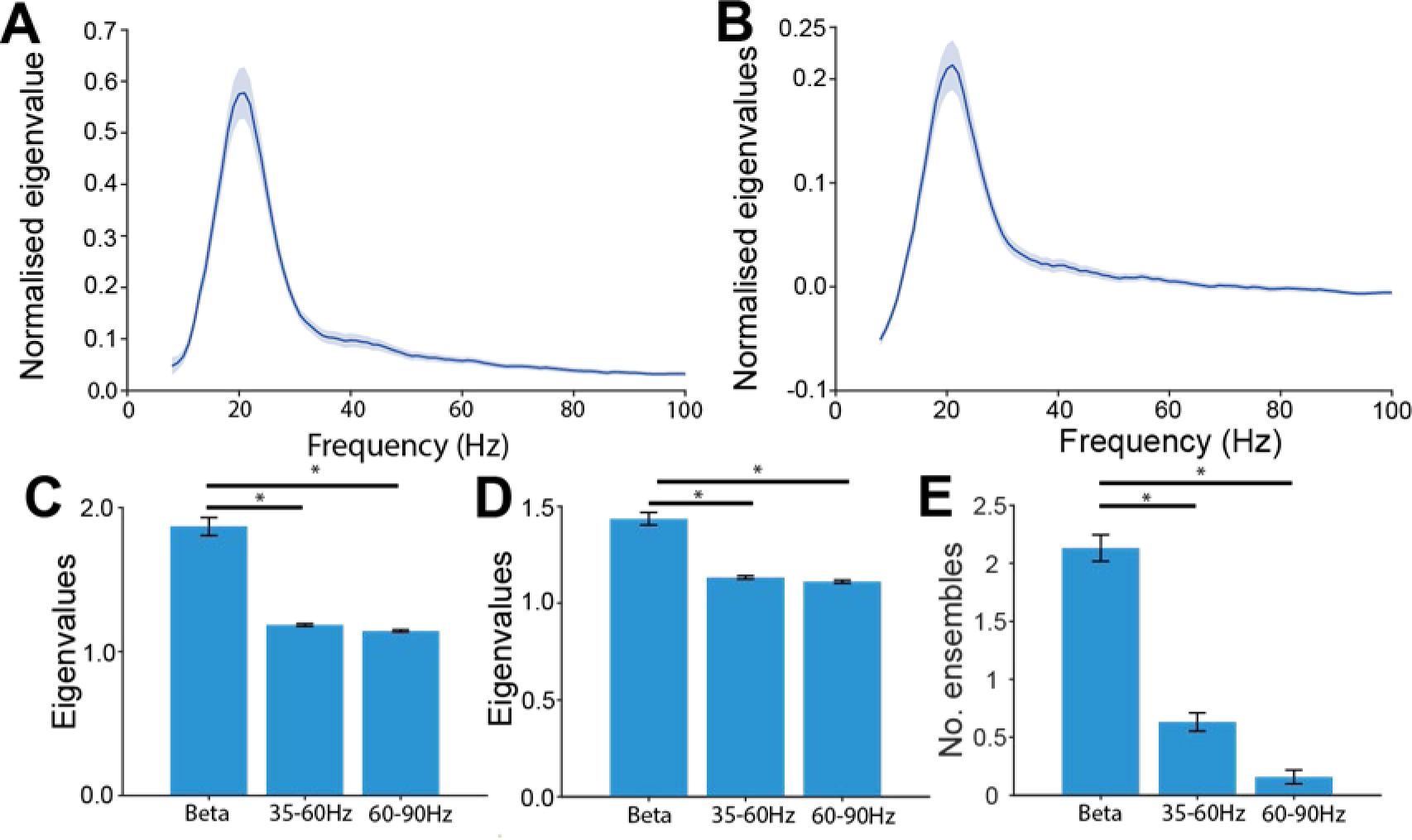
Coordinated oscillation in BUA was specific to the beta frequency band: **A** and **B:** The largest (**A**) and the average of the 3 largest (**B**) normalized eigenvalues of the correlation matrix for the change in envelope of BUA filtered at ±5Hz of the centre frequency. **C, D** and **E**: The change in envelope of beta oscillations (15-25Hz) was compared to low (35-60 Hz) and high (60-90 Hz) gamma in terms of the size of the first eigenvalue (**C**, Wilcoxon sign rank test, beta-low gamma: p<10^-7^, beta-high gamma: p<10^-7^), the average of the first 3 eigenvalues (**D,** Wilcoxon sign rank test, beta-low gamma: p<10^-7^, beta-high gamma: p<10^-7^), and the number of ensembles identified (**E**, Sign test, beta-low gamma: p<10^-8^, beta-high gamma: p<10^-10^).

### Spatially coordinated beta emergence across time

We next sought to understand the changes in instantaneous beta amplitude and phase sychrony across space during spatially coordinateed beta emergence and offset. The expression strength of coactivity patterns was computed (**Fig 7A**). Peaks in expression strength (beta ensemble activations) were primarily driven by large co-ordinated changes in the envelope of beta-BUA in at least a pair of channels with high contributions to the beta ensemble (**Fig 7B-C**). The low stable baseline punctuated by large peaks reflected that much of the coordinated change in beta envelopes occurred as brief, transient events. Whilst these activation events were solely identified on the basis of changes in envelope, they were also accompanied by large changes in phase synchrony between member channels (**Fig 7D-E**).

**Figure 7.**
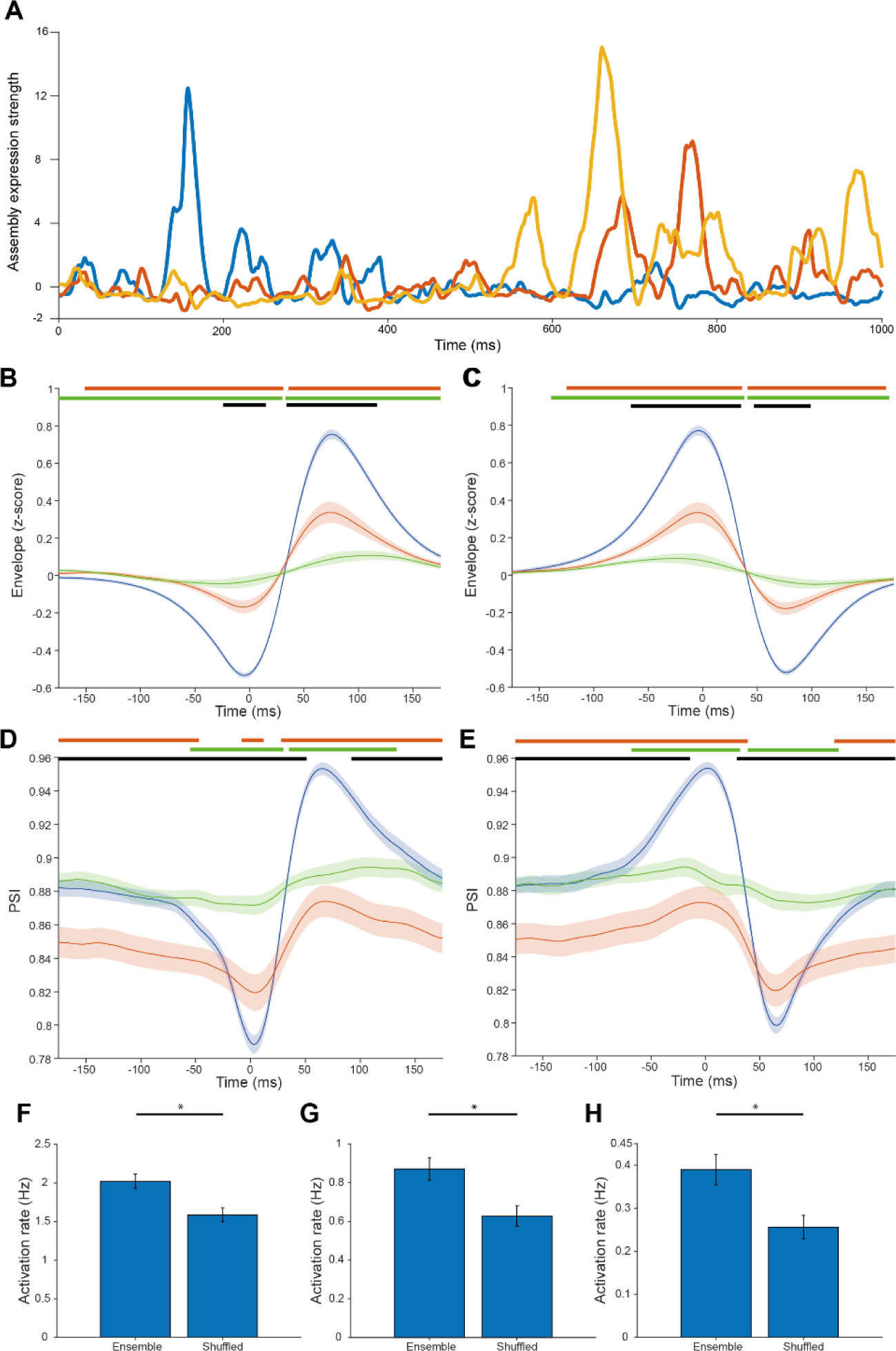
coactive changes in beta envelope in beta ensembles were associated with increases in phase synchrony specifically in member channels: **A:** The expression strength of the coactivity patterns of 3 ensembles (identified in a single recording) over time. **B** and **C**: The average beta-envelope triggered by ensemble activations (activation threshold = 5), separating activations by whether they were associated with a coordinated increase (**B**) or decrease (**C**) in beta envelope. These were associated with a larger coordinated increase and decrease in beta-envelope for **B** and **C** respectively in member channels of the activating (blue) than non-activating (green) ensembles or non-member channels (orange). **D** and **E**: The average PSI triggered by ensemble activations (activation threshold = 5), separating activations by whether they were associated with a coordinated increase (**D**) or decrease (**E**) in beta envelope. These were associated with a larger coordinated increase and decrease in PSI for **D** and **E** respectively in member channels of the activating (blue) than non-activating (green) ensembles or non-member channels (orange). For figures **B**, **C**, **D** and **E** periods of significant difference are marked above the graph. The orange marker signifies periods of significant difference between members of the activating ensemble and non-member channels. The green marker signifies periods of significant differences between members of activating and non-activating ensembles The black line signifies periods of significant difference between the members of non-activating ensembles and channels that are not members of any ensemble. Significance was determined with the Wilcoxon rank sum test using false discovery rate statistics to control for the multiple time points compared. **F**, **G** and **H**: The frequency of activations was greater in ensembles compared with circularly shifted ensembles, with an activation threshold of 5 (Wilcoxon signed-rank test, p= 1.7 x 10^-10^), 7.5 (Wilcoxon signed-rank test, p= 8.8 x 10^-11^) and 10 (Wilcoxon signed-rank test, p= 2.5 x 10^-10^) respectively.

These activations were significantly more frequent in true beta ensembles than in pseudo-beta ensembles formed by circularly shifting the assembly pattern for each beta ensemble by a random amount (**Fig 7F-H**). This difference in activation rate between beta ensembles and pseudo-beta ensembles became increasingly pronounced as the threshold of activation was increased. Taken together, these findings confirm that coactivity in the emergence of beta oscillations is specific to the identified beta ensembles and is accompanied by an increase in phase synchrony.

### Subcortical ensembles and their relationship with respect to ECoG

Cortical beta-bursts have been associated with increases in beta envelope in the GP and STN and increased phased synchrony between basal ganglia structures and the cortex (Cagnan et al. 2019). Previous work has probed the relationship between ECoG and the basal ganglia at a brain structure-by-structure level. We aimed to test whether beta neural populations in the GP that showed coordinated beta oscillation emergence had a different relationship with cortical beta oscillations to non-members. Both member and non-member channels had a large rise in their coherence with ECoG at ∼15-30 Hz, which peaked at ∼20.5 Hz. This peak in coherence was larger with BUA channels that were ensemble members (**Fig 8A**). Ensemble activations were divided by whether they represented a coordinated increase or decrease in the envelope of beta-BUA. Ensemble activations were associated with moderate changes in the envelope of beta-ECoG (**Fig 8B, C**). This moderate amplitude likely reflects that some, but not all, ensemble activations are associated with beta bursts in the cortex. Equally, the change in PSI across ensemble activations between beta-ECoG and beta-BUA in member channels was greater than for channels that were not members of the activating beta ensemble (**Fig 8D, E)**. Overall, membership of a beta ensemble predicted that a given channel was more strongly related to cortical beta.

**Figure 8.**
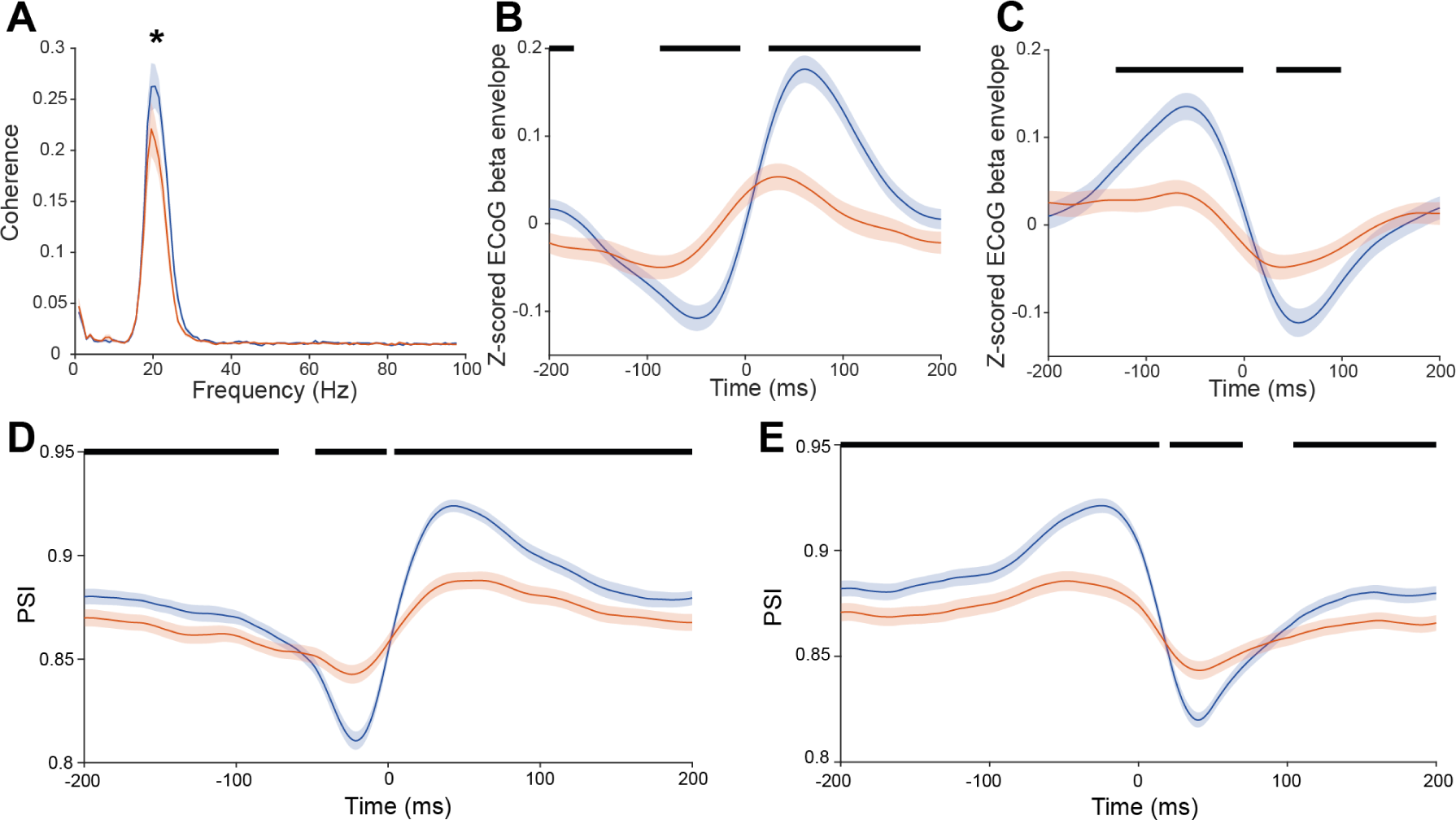
Beta ensemble activations were associated with changes in both the instantaneous power of cortical beta and the phase synchrony in the beta-band between the beta ensemble and the cortex: **A**: The peak coherence of the BUA of member channels (blue) with ECoG was greater than that of non-member channels (orange) (Wilcoxon rank sum test, p<0.05). **B** and **C:** Average beta-ECoG envelope triggered by ensemble activations (blue) separating by whether these activations reflected a coordinated increase (**B**) or decrease (**C**) in the envelope of beta-BUA respectively. Also plotted was the triggered average of beta-ECoG envelope at random time points (orange) separating for whether these random time points correspond to a coordinated increase (**B**) or decrease (**C**) in the envelope of beta-BUA in the beta ensemble. **D** and **E:** PSI between beta-ECoG and beta-BUA averaged across ensemble activations in member (blue) and non-member (orange) channels, separating by whether these activations reflected a coordinated increase (**D**) or decrease (**E**) in the envelope of beta-BUA respectively. **B**, **C**, **D** and **E** have periods of significant difference between traces displayed with a black marker. Significance was determined with the Wilcoxon rank sum test using false discovery rate statistics to control for the multiple time points compared.

**Figure 9:**
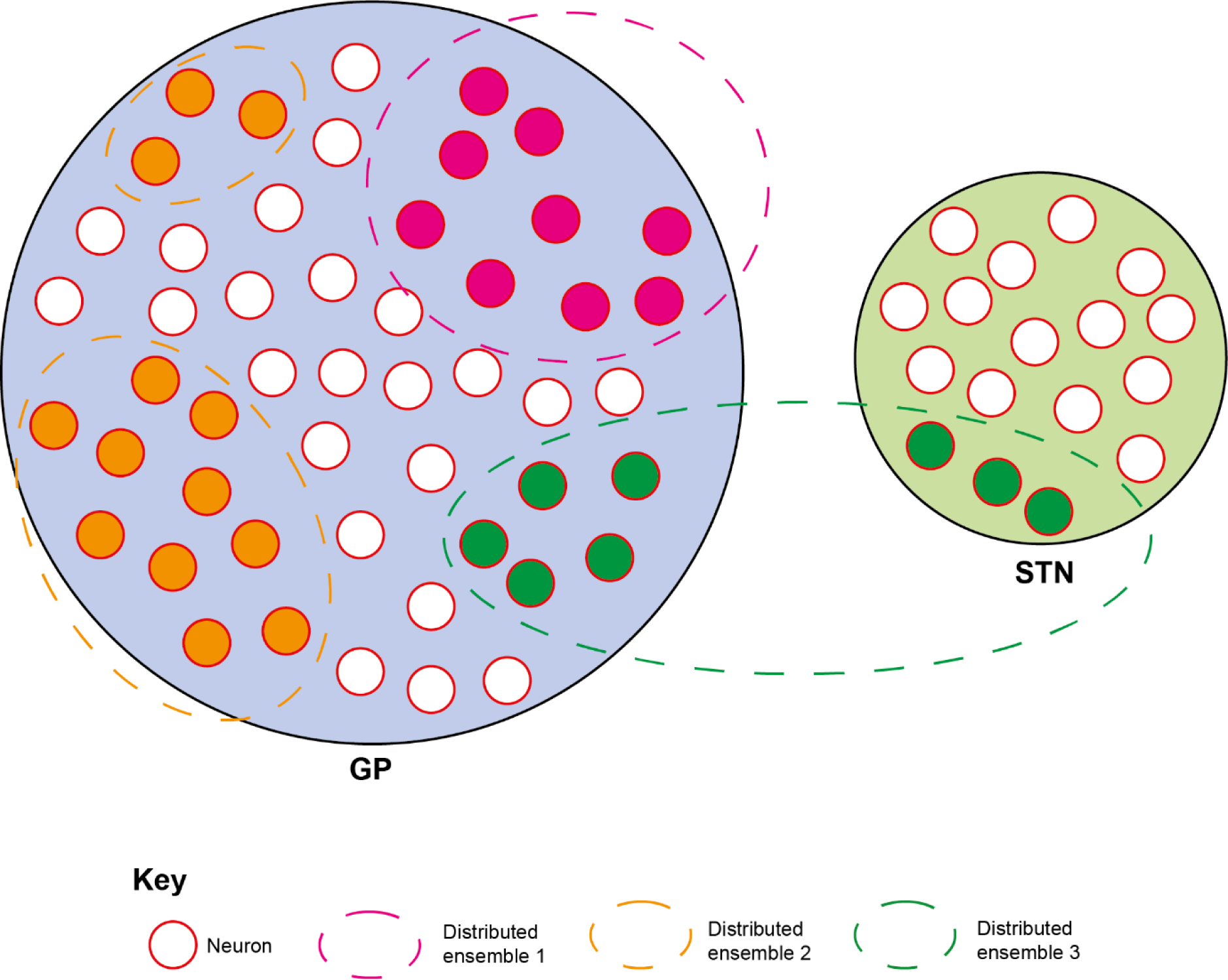
Neurons across the GP and STN form distinct beta ensembles which each have coordinated changes in the envelope of their beta oscillations. Beta oscillations evolve synchronously across the neural population that makes up a beta ensemble, which can form within or across anatomical structures. As coordinated beta activity is accompanied by an increase in beta phase synchrony in the neurons that make up beta ensembles, they likely indicate dynamic synchronisation of spike timing across large spatial extents of the basal ganglia.

## Discussion

Abnormally sustained, transient increases in the power of beta oscillations are markers of impaired network activity in the parkinsonian brain. How these transient increases in beta synchrony manifest on the spatial scale, however, is less clear. Here we provide a novel application of a well-established analytical technique to define the nature of spatial synchronization in multichannel unit recordings from parkinsonian rats. The results demonstrate that the spiking of multiple, independent ensembles of neurons can be distinctly and transiently synchronized by beta oscillations within the cortico-basal ganglia network. These findings have potentially important implications for the interpretation of beta oscillations as a biomarker of pathophysiology in PD and its treatment with DBS.

### Anatomical properties of cortico-basal ganglia circuits and beta oscillations

An influential hypothesis as to the pathophysiological mechanism in PD is that the functional segregation of cortico-basal ganglia loops is compromised (Filion et al., 1988; Tremblay et al., 1989; Boraud et al., 2000; Leblois et al., 2006), preventing the normal function of those circuits. Cortico-basal ganglia circuits are arranged into information streams that can be broadly divided by cortical inputs to the striatum (Alexander et al., 1990). These were traditionally thought to consist of 3/4 functional subdivisions, but recent studies have shown that in rodents there are 10s-100s of cortico-basal ganglia-thalamic loops in the mouse brain (Hunnicutt et al., 2016; Peters et al., 2021). Inputs from functionally related cortical areas converge into specific parts of the striatum, but these clusters remain segregated in their axonal projections to the pallidum (Foster et al., 2021). However, there is significant convergence of these striatopallidal channels onto the basal ganglia output nuclei and the subsequent thalamocortical projection. Overall, cortico-basal ganglia circuits are, therefore, partially segregated with multiple points of convergence. The expression and propagation of beta oscillations will be partly defined by these anatomical features. As previously suggested, the spatial hypersynchronization of beta oscillations could bind the activity of neurons across usually segregated loops, constraining the computation space (Brittain et al., 2014; Cagnan et al., 2015). To our knowledge, however, the spatial extent of such hypersynchronization has not been systematically quantified. By providing an approach to quantify the dependence of beta oscillations across space, we aimed to provide novel insights into this issue.

### Defining spatial synchronization of beta oscillations at the neuronal level

We analysed the temporal correlation of the change in beta amplitude of BUA signals recorded from equally spaced recording channels on silicon probes. We chose the power of the BUA as our primary signal for analysis because it corresponds to the level of beta synchronization of spiking in a small pool of neurons around the electrode (Sharott et al., 2017; Cagnan et al., 2019). This is a useful signal given that transient increases in beta synchronization appear to be a key pathophysiological feature of the disease (Tinkhauser et al., 2017a; Cagnan et al., 2019; Baaske et al., 2020). When the fluctuations in BUA beta amplitude are positively correlated across electrode channels, it suggests that the same process is coordinating local synchronization across those areas. For analysis of such temporal correlation to be meaningful, it is important that it results from neuronal processes, rather than passive ones such as volume conduction. LFPs are therefore suboptimal for this purpose, as re-referencing cannot guarantee independence and more complex signal processing complicates interpretation. In contrast, the change in envelope of beta-BUA signals have a high level of independence without further referencing, with even adjacent channels outside of the beta ensembles explaining an average of only 1% of each other’s variance. Thus, when fluctuations in the envelope of beta-BUA are significantly correlated, it is likely to be driven by neuronal activity.

To quantify these correlations across every channel in a given recording, we employed a PCA/ICA framework that is well-established for the study of single units (Peyrache et al., 2010; Lopes-dos-Santos et al., 2011; van de Ven et al., 2016; Lopes-Dos-Santos et al., 2018). The key feature of this method is that it defines *groups* of channels with correlated changes in the amplitude of beta oscillations in their spiking activity that are otherwise independent of each other and is agnostic to distance. Interestingly, the expression strength of assembly patterns tended to have a stable baseline with transient peaks. This mirrors findings from ensembles identified in single units, where again coordinated firing is dominated by short duration events (Lopes-dos-Santos et al., 2013). These peaks likely represent coordinated beta bursts across beta ensembles. In the majority of recordings, this method detected more than one ensemble. This result suggests that beta synchronization is not a homogenous process across circuits, but there are multiple, distinct beta “activities” that can couple across specific neuronal populations independently of each other. In our experimental set-up, we cannot identify the substrate for such independent beta activities, but it seems likely that they could be the result of the anatomical segregation of basal ganglia circuits described above. Importantly, many ensembles were not spatially contiguous and/or spanned the pallidum and STN. Ensembles may therefore have represented pools of neurons connected by common inputs, rather than simply being in direct proximity to each other.

### Physiological properties of beta ensembles

The ensembles defined by our analysis were based on fluctuations in beta amplitude, but were accompanied by enhanced beta phase synchronization with other ensemble members and the cortex. This beta synchronization is likely to be the cause, rather than effect of coordinated fluctuations in beta power across channels. We have already described the transient increase in phase locking of individual units/BUA channels during cortical beta bursts (Cagnan et al., 2019). Here, we extend those observations to show that cortical beta bursts are accompanied by transient beta synchronization across spatially distinct populations of basal ganglia neurons. In addition, the raw BUAs (i.e. not beta filtered) of member channels were around 100% more correlated with each other at zero lag than with non-members. This indicates that groups of neurons that were spatially synchronized on the scale of beta oscillations were also far more likely to fire synchronous spikes within a few milliseconds of each other (Buzsaki, 2010). This finding is important, as it indicates that ensembles will produce highly synchronized outputs that are likely to propagate to, and temporally summate within, downstream targets. More speculatively, correlations on these time scales could contribute to maladaptive plasticity in the parkinsonian brain (Chu et al., 2015). On a technical level, using phase-synchronisation or cross correlation between channels to define ensembles may have led to similar results as using beta amplitude. However, changes in beta power provides a more straightforward signal for ICA/PCA and can readily be applied in other contexts for future studies. More importantly, our method provided an effective way of identifying spatially distributed channels that shared features relating to oscillation and synchronization.

One potential caveat of our findings is the inability to fully distinguish two possible mechanisms through which the coordination reported here could arise. Mechanism 1: Beta ensembles reflect the coordinated emergence of beta oscillation in spiking activity of spatial extents of the basal ganglia made up of neurons proximal to each of the recording contacts. This is the interpretation we think is most likely. Mechanism 2: That beta ensembles are driven by large oscillations in the spiking output of focal neural ensembles that are detectable across multiple channels. In mechanism 2, PCA-ICA could be thought of as identifying ‘sources’ of beta oscillations. In this interpretation, several such distinct ‘sources’ can be identified within and across anatomical structures. Therefore, regardless which of the two mechanisms outlined above are at play, this paper provides evidence that there are neural populations within the basal ganglia with distinct emergence to their beta oscillations, be this spatial extents of the basal ganglia or more focal neural populations with oscillations large enough to be recorded across multiple contacts. In our opinion, Mechanism 1 remains the more likely of the two interpretations, as members of beta assemblies are more correlated even after controlling for distance. These correlations were seen up to 400 microns, a distance where it is highly unlikely the same spiking activity can be recorded. Assembly patterns identified could also be spatially non-contiguous, span recording probes and even anatomical structures.

### Relevance to PD pathophysiology and DBS

As the recordings used here were made in anaesthetized rats, we cannot guarantee the generalisability of our findings to the awake state. Nevertheless, the principles identified here raise specific hypotheses as to how spatial synchronization of beta activities are related to Parkinsonian symptoms. Beta oscillations, quantified in a variety of ways, are most strongly related to akinetic/rigid symptoms (Kühn et al., 2005; Sharott et al., 2014; Neumann et al., 2016). Within each patient, the severity of akinetic rigid symptoms often varies between upper and lower body and individual limbs. Our findings suggest that independently synchronized ensembles could underlie this variance and provide a mechanism through which different limbs could become bradykinetic at a given time.

Adaptive DBS targeting beta bursts has been found to outperform continuous stimulation in patients with Parkinson’s disease (Little et al., 2013). This allows large reductions in the electrical energy that must be delivered to control symptoms, while reducing the rate of battery depletion and the severity of side effects (Little et al., 2013). Stimulation is typically delivered with a single stimulating electrode within a single brain structure (i.e., STN or GP, not both). Given that spatially clustered subregions in the basal ganglia show coordinated changes in the envelope of beta activity, there may also be value to spatial targeting for adaptive DBS protocols. Such a stimulation strategy was pioneered by Tass and colleagues, where electrical stimulation has been delivered from different contacts of a stimulating electrode in an open loop fashion to reset the coordinated neural activity observed in Parkinsonism (Tass, 2003; Tass et al., 2012). The main advantages of coordinated reset over other adaptive stimulation approaches are long term plastic changes and delayed symptom return (Tass et al., 2012). As discussed above, ensembles in our data were synchronized at timescales relevant for the induction of plasticity. More targeted interference of such spatiotemporal synchronization thus has the potential to interfere with this process and could result in larger long-term effects on symptom expression. The use of segmented leads could potentially facilitate such an approach (Debove et al., 2023). Future work may focus on combining both adaptive stimulation approaches and deliver targeted bursts of stimulation to spatiotemporally distinct pockets of ensemble activity to further reinforce the desynchronizing effects of stimulation and potentially induce longer lasting plastic changes.

## Data Availability Statement

The preprocessed data is available at: https://data.mrc.ox.ac.uk/data-set/wideband recordings-silicon-probes-subthalamic-nucleus-6-ohda-hemi-lesioned-rats-during and has DOI: 10.5287/bodleian:wx6D7oenk

