## Supplemental Figure 1. for "Beta bursts in the parkinsonian cortico-basal ganglia network form spatially discrete ensembles"

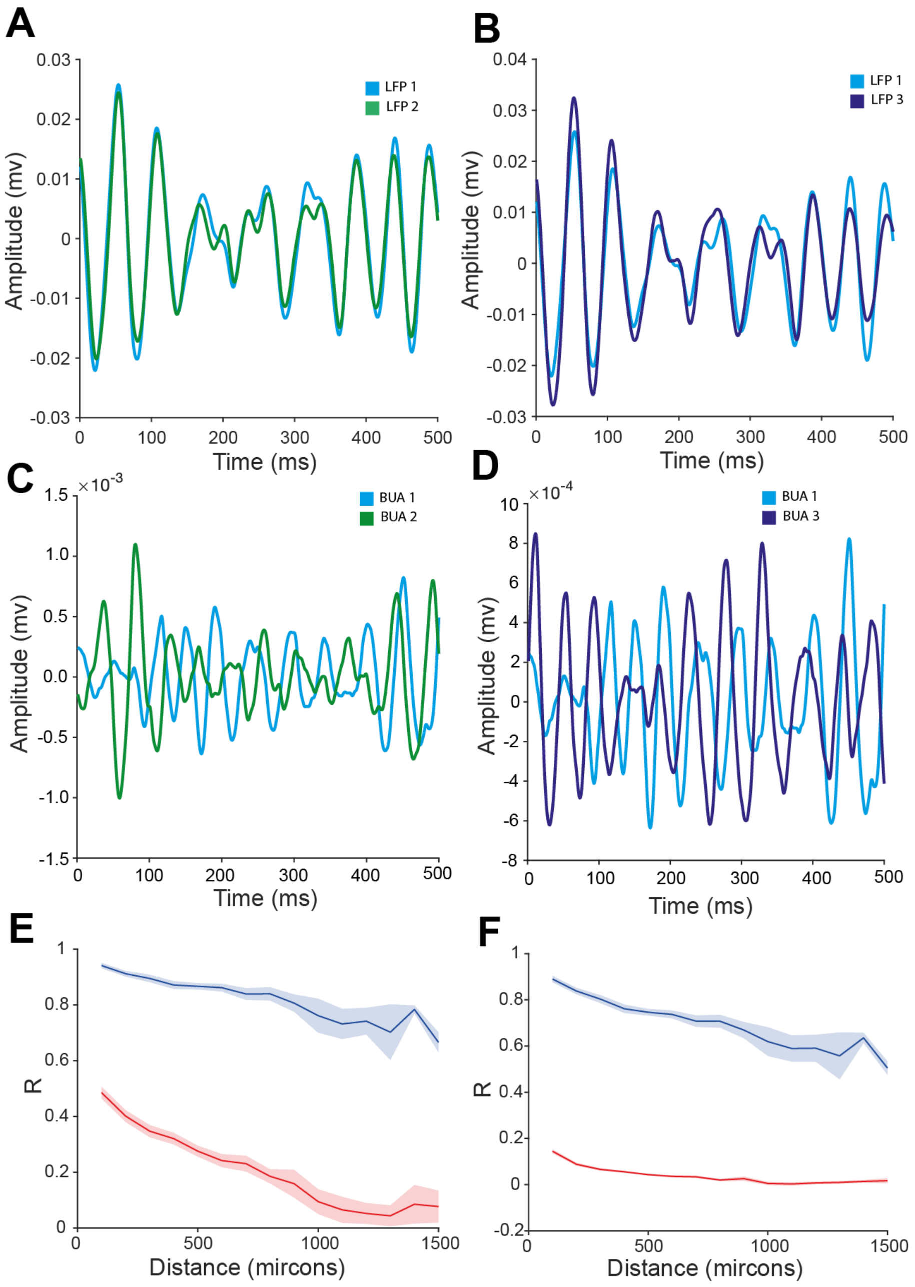


**Figure S1, correlations in the change of the envelope of beta filtered BUA do not arise as a result of volume conduction**: **A** and **B**: Exemplary beta filtered LFPs from 2 channels separated by 100 microns (LFP 1 and LFP 2 coloured in blue and green respectively) and 1000 microns (LFP 1 and LFP 3 coloured in blue and purple respectively) respectively. **C** and **D**: Exemplary beta filtered BUAs from 2 channels separated by 100 microns (BUA 1 and BUA 2 coloured in blue and green respectively) and 1000 microns respectively (BUA 1 and BUA 3 coloured in blue and purple respectively). **E**: The average absolute Pearson's R of the beta-filtered LFP (blue) or beta-filtered BUA (red) between pairs of channels over variable distances. **F**: The average absolute Pearson's R of the change in beta-envelope over 50ms of the LFP (blue) or BUA (red) between pairs of channels over variable distances. The correlation between the beta-filtered signal and the change in the beta envelope over 50ms was much greater across pairs of LFPs than BUAs across all distances. Furthermore, whilst the correlation across all measures decreased with distance, this decrease was less steep for the LFP-derived signals than for BUAs.
